# Pattern of severe injuries in Spanish children: boys and falls are alarmingly overrepresented

**DOI:** 10.1101/303958

**Authors:** Sergio Benito, Rosa López-Parellada, Elisabeth Esteban, Lluïsa Hemández-Platero, Salvi Prat, Mireia Esparza, Francisco José Cambra, Maria Esther Esteban

**Affiliations:** Pediatric Intensive Care Unit Service, Hospital Sant Joan de Déu. Passeig Sant Joan de Déu 2, 08950 Esplugues de Llobregat, Barcelona, Spain.; Section of Zoology and Biological Anthropology, Department of Evolutionary Biology, Ecology and Environmental Sciences, Faculty of Biology, Universitat de Barcelona, Avda. Diagonal 643, 08028 Barcelona, Spain.; TraumCat coordinator, Hospital Clínic Barcelona, Carrer de Villarroel, 170, 08036 Barcelona, Spain.; Institut de Recerca de la Biodiversitat (IRBio), Universitat de Barcelona, Avda. Diagonal 643, 08028 Barcelona, Spain.

**Keywords:** multiple injury, trauma systems, fall, descriptive epidemiology, child

## Abstract

**Background**: Taking into account that injury is one of the main causes of child fatalities in developed countries, and that boys are more likely to suffer it than girls, we have explored a database of pediatric patients with severe injuries to determine whether sex and age influence the pattern of these fatalities, and the magnitude of this.

**Method**: Observational study of the demographic and clinical characteristics of 227 patients from a Spanish pediatric reference hospital, all of them admitted with a diagnosis of trauma.

**Result**: Falls are the most frequent type of trauma (60.7%), followed by pedestrian traffic collisions (15%). Boys are over-represented in falls (72% *vs* 28% in girls) and pedestrian traffic injuries (61% *vs* 39 %). In boys, falls are mainly observed in public roads and during leisure activities (53.8%) whereas in girls at home (55.2%). In a logistic regression, sex and age are statistically significant predictors of severe trauma, boys (OR = 1.59) and the adolescent age group (OR = 3.7) showed the highest odds.

**Conclusion**: We have observed a clear gender-biased pattern of injury-related events: falls are the leading cause of injuries, with 2.5 boys for every girl. Falls mostly happened during outdoor leisure activities in boys and at home in girls. Pedestrian traffic injuries also show significant differences between sexes, emphasizing the role of cognitive and cultural factors in children’s behavior.

## WHAT IS ALREADY KNOWN ON THE SUBJECT

- In 2016 road injuries were the third cause of death among children from 5 to 9 years old in the OCDE countries.
- Road traffic injuries and falls are the most relevant sources of unintentional trauma.
- Injury-related events are gender-biased, with boys suffering more frequent and severe events than girls. But there are few studies indicating whether this sex biased pattern is present in all types of trauma.

## WHAT THIS STUDY ADDS

- Sex and age are statistically significant predictors of severe trauma in children. Boys show 2.5 times more severe falls than girls, mainly during outdoor leisure activities.
- Cognitive and cultural factors may explain the remarkable age differences observed between boys and girls in pedestrian traffic collisions.
- The relatively high proportion of young girls involved in falls from great heights needs to be contrasted with data from comparable studies, and if this tendency is confirmed, preventive measures must be launched.

## INTRODUCTION

Although every region in the world has reduced its child mortality rate, there are still large differences among rich and poor nations. Infectious diseases and neonatal complications are the main killers of children from poor nations while injury is now one of the leading killers of children in developed countries. In 2001, the *UNICEF’s Innocenti Report Card* was devoted to the analysis of child deaths by injury in rich nations [1]. At that time, injury accounted for 40% of deaths in children from 1 to 14 years in the OECD (Organisation for Economic Cooperation and Development) countries. According to more recent data, in 2016 road injuries were the third cause of death among children from 5 to 9 years old [2]. Road traffic injuries and falls are the most relevant sources of unintentional trauma and the main cause of traumatic brain injury (TBI), especially in young children [3–5]. For instance, in Spain, the country where this investigation has been conducted, 66% of severe brain injuries in children below the age of 2 years were due to falls [6].

As we stated above, trauma is a relevant public health issue leading to elevated mortality, especially in children. Severe trauma patients are characterized by having multiple injuries produced in a single event in which at least one potentially implies a vital risk. Children show different patterns of injury from adults because of their anatomy, behavior, and their changing levels of maturity. Head trauma combined with fractures is the most frequent pattern in pediatric multiple trauma patients, followed by a combination of head and chest or abdominal trauma [7,8].

Many studies have evaluated the effects of severe trauma within the general population without making a distinction between adults and children, with only a few focusing exclusively on pediatric multiple trauma [9–12]. These studies are basically descriptive including but not limited to information on the mechanism of trauma, the trauma scoring systems and patients’ physiological parameters [13,14]. In 2015 we analyzed a large sample of children admitted in a pediatric intensive care unit to explore the role of severe stress on sex-related survival. Although girls seemed to be more vulnerable because of a wider range of hospital admission causes, the proportion of boys exceeded the proportion of girls in admissions due to multiple trauma [15]).

It is not new that injury-related events are gender-biased, with boys suffering more frequent and severe injuries than girls. Differential exposure to risky activities, higher activity levels and less careful behavior may contribute to explain this fact [16,17]. In this work we have analyzed a database of children admitted in a reference hospital due to severe trauma to determine whether this sex biased pattern is present in all types of trauma, and the magnitude of it. To do that we have explored the type and place of trauma event, and the demographic and clinical characteristics of the children involved in it. These data will help us to determine to what extent sex and age are determinants of severe injuries in children, and hence, to launch the necessary awareness measures.

## MATERIALS AND METHODS

This observational study has been conducted with the available data of 227 patients admitted to the Hospital Sant Joan de Déu (Esplugues de Llobregat, Barcelona, Spain) with a diagnosis of severe trauma. Eighty-three percent of trauma events were located in the province of Barcelona, 61% in the metropolitan area and the rest in different localities of the Autonomous Community of Catalonia. Sant Joan de Déu is a pediatric public hospital of reference for the whole Autonomous Community (1,185,410 inhabitants in the age group of 0-14 years; Institut d’Estadística de Catalunya IDESCAT).

The data included in this study have been obtained from a Trauma Registry (Registro de Traumatismo Grave en Cataluña, TraumCat) created in Catalonia in 2011 to guide and control the care process of severe trauma [18]. This platform, under the Public Health Service of Catalonia, collects hospital based information from trauma patients allowing the evaluation of the trauma care process between the prehospital care and the accredited trauma care centers for complete description see Prat *et al*. [19]. The coordination between the medical emergency service and the hospitals is done through a polytrauma patient code (PPT code). This code is activated by the medical emergency service from the injury scene. Once patients with severe trauma are *in situ* attended and stabilized, the PPT code maintains a high level of care during transport, reception at the destination hospital and transfer, reducing care time, diagnosis and treatment. In children, the PPT code is activated in response to vital signs, anatomy of the injury, type of trauma and medical history. There are four types of criteria for PPT code activation (ranging from PPT 0 to PPT 3) based on physiological, anatomical and biomechanical characteristics as well as comorbidity or other considerations [19].

All trauma patients admitted to the pediatric intensive care unit (PICU) and all patients admitted to the hospital with a PPT code of 0, 1 or 2 between July 2012 and January 2016 were considered for the study. Data from each patient came from the first data recorded by the medical emergency service, from the hospital reception, and from the patients’ medical history. Injuries and mechanisms of injury were registered according to the International Classification of Diseases, Ninth Revision, Clinical Modification (ICD-9-CM) [20].

The main variables included in the study were: age, sex, type and origin of trauma, distribution of events according to the month and day of the week, trauma location and type of vehicle involved in traffic collisions. According to the World Health Organization age was grouped as a categorical parameter as follows, neonate (0 to 30 days of age), infant (1 month to 2 years), young child (2 years to 6 years), child (6 years to 12 years) and adolescent (12 years to 18 years) [21]. To estimate the severity of injury, the database recorded the affected body region and standardized scoring systems such as the Glasgow Coma Scale (GCS) [22] and the Pediatric Trauma Score (PTS) [23].

Data were analyzed using the IBM^®^ SPSS^®^ Statistics 23.0 software (IBM^®^ Corporation. Armonk, NY). Qualitative variables were described through frequencies and percentages, whereas quantitative variables were described through measures of central tendency. The Kolmogorov-Smirnov test was used to assess normality of quantitative data. Parametric variables were compared with Student’s t test. Nonparametric variables were compared with Mann-Whitney U test. Fisher exact test and Chi-square test were used to evaluate differences between qualitative variables. A p value below 0.05 was considered statistically significant.

After the univariate analysis of data, a binary logistic forward regression was performed to establish whether sex and age were risk factors for severe trauma. To carry out this analysis, the database of pediatric patients of Esteban *et al*. [15] was used. Girls and neonates were used as the reference group. Results were presented as odds ratios (ORs) with 95% confidence intervals (CI).

## RESULTS

The descriptive characteristics of the sample are shown in Table 1. We found significantly (p = 0.001) more boys than girls admitted (66.5% *vs* 33.5%). The median age was 6.5 years in boys and girls. As expected, neonates and infants were the categories with the lowest number of cases. According to the origin of trauma, 90.2% of events occurred unintentionally and only 5.8% were caused intentionally. Within the category of intentional injuries, four (33.3%) were self-harm, two (16.7%) were aggressions, three (25%) were child abuse and three cases were classified in other categories. The distribution of trauma by type was similar in both sexes: falls were the predominant trauma event (65.7% in boys, 50.7% in girls), followed by a distance by being run over (13.9% boys, 17.4% girls) and traffic collisions (9.5% boys, 15.9% girls). The trauma site showed remarkable sex differences (p = 0.002): girls suffered substantially more injuries at home (41.9% girls *vs* 25.4 % boys) whereas boys had more injuries during leisure activities (21.4% boys *vs* 4.8% girls).

With regard to the seasonal distribution of trauma events, although the differences were not statistically significant, girls had more events in summer (45.3%) while boys, with the exception of winter, showed a quite similar number of events during the rest of the year (from 28.1% in spring to 32.3% in summer). When we looked at the daily distribution of injuries we could not attribute a greater number of them to any specific day of the week regardless of the season (data not shown).

**Table 1.**
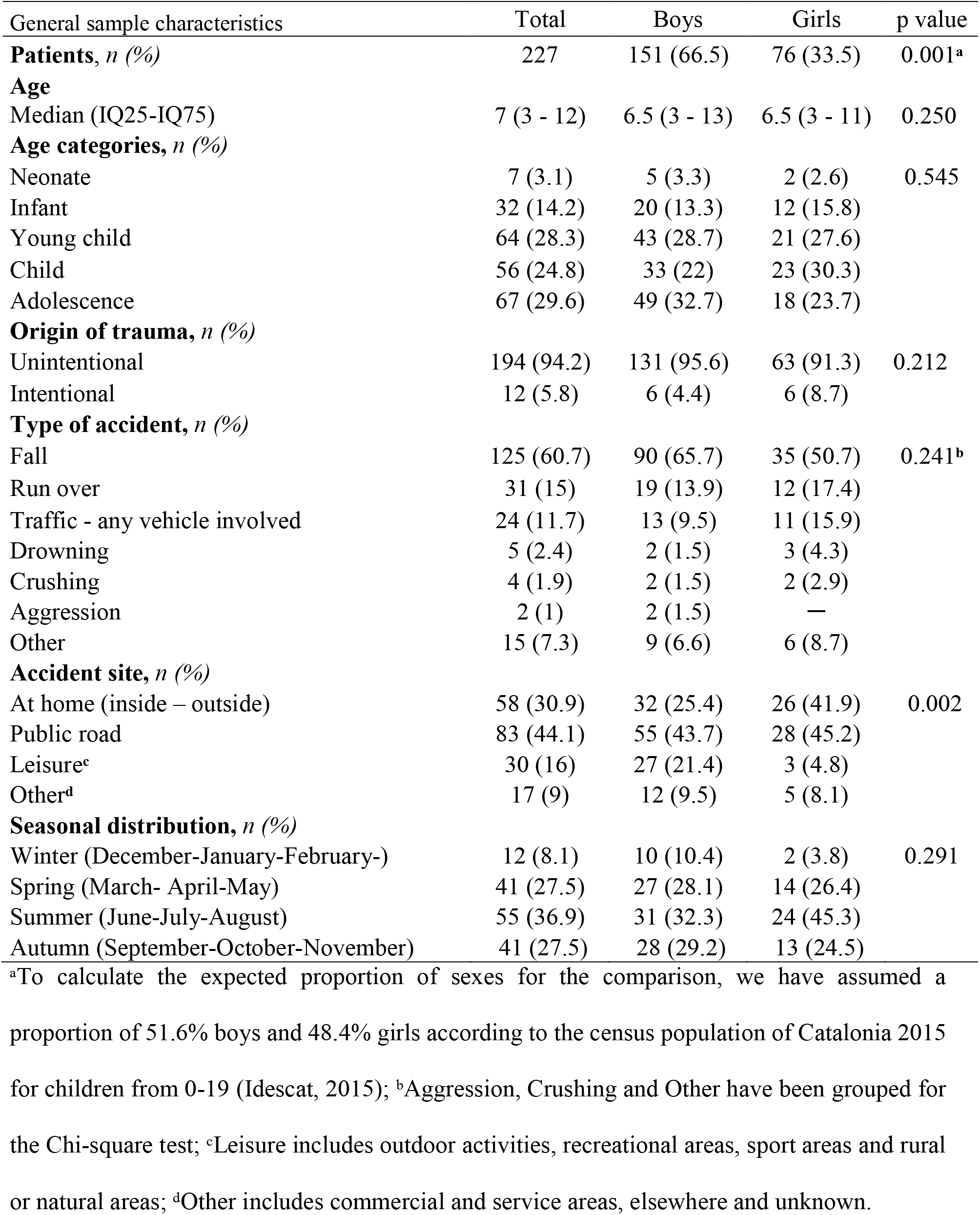
Demographic characteristics of the sample. P value indicates the results of sex comparisons.

| General sample characteristics | Total | Boys | Girls | p value |
| --- | --- | --- | --- | --- |
| **Patients, *n* (%)** | 227 | 151 (66.5) | 76 (33.5) | 0.001<sup>a</sup> |
| **Age** |  |  |  |  |
| Median (IQ25-IQ75) | 7 (3 - 12) | 6.5 (3 - 13) | 6.5 (3 - 11) | 0.250 |
| **Age categories, *n* (%)** |  |  |  |  |
| Neonate | 7 (3.1) | 5 (3.3) | 2 (2.6) | 0.545 |
| Infant | 32 (14.2) | 20 (13.3) | 12 (15.8) |  |
| Young child | 64 (28.3) | 43 (28.7) | 21 (27.6) |  |
| Child | 56 (24.8) | 33 (22) | 23 (30.3) |  |
| Adolescence | 67 (29.6) | 49 (32.7) | 18 (23.7) |  |
| **Origin of trauma, *n* (%)** |  |  |  |  |
| Unintentional | 194 (94.2) | 131 (95.6) | 63 (91.3) | 0.212 |
| Intentional | 12 (5.8) | 6 (4.4) | 6 (8.7) |  |
| **Type of accident, *n* (%)** |  |  |  |  |
| Fall | 125 (60.7) | 90 (65.7) | 35 (50.7) | 0.241<sup>b</sup> |
| Run over | 31 (15) | 19 (13.9) | 12 (17.4) |  |
| Traffic - any vehicle involved | 24 (11.7) | 13 (9.5) | 11 (15.9) |  |
| Drowning | 5 (2.4) | 2 (1.5) | 3 (4.3) |  |
| Crushing | 4 (1.9) | 2 (1.5) | 2 (2.9) |  |
| Aggression | 2 (1) | 2 (1.5) | — |  |
| Other | 15 (7.3) | 9 (6.6) | 6 (8.7) |  |
| **Accident site, *n* (%)** |  |  |  |  |
| At home (inside – outside) | 58 (30.9) | 32 (25.4) | 26 (41.9) | 0.002 |
| Public road | 83 (44.1) | 55 (43.7) | 28 (45.2) |  |
| Leisure<sup>c</sup> | 30 (16) | 27 (21.4) | 3 (4.8) |  |
| Other<sup>d</sup> | 17 (9) | 12 (9.5) | 5 (8.1) |  |
| **Seasonal distribution, *n* (%)** |  |  |  |  |
| Winter (December-January-February-) | 12 (8.1) | 10 (10.4) | 2 (3.8) | 0.291 |
| Spring (March- April-May) | 41 (27.5) | 27 (28.1) | 14 (26.4) |  |
| Summer (June-July-August) | 55 (36.9) | 31 (32.3) | 24 (45.3) |  |
| Autumn (September-October-November) | 41 (27.5) | 28 (29.2) | 13 (24.5) |  |

<sup>a</sup>To calculate the expected proportion of sexes for the comparison, we have assumed a proportion of 51.6% boys and 48.4% girls according to the census population of Catalonia 2015 for children from 0-19 (Idescat, 2015); <sup>b</sup>Aggression, Crushing and Other have been grouped for the Chi-square test; <sup>c</sup>Leisure includes outdoor activities, recreational areas, sport areas and rural or natural areas; <sup>d</sup>Other includes commercial and service areas, elsewhere and unknown.

Sex differences in falls, the most common cause of trauma, are detailed in Table 2. Boys largely exceeded the number of girls (72% *vs* 28%; p = 0.038). The median age is 8 years old in both sexes. Concerning the trauma site, and consistent with that observed for the whole data, boys suffered significantly (p = 0.0002) more events during leisure activities (27.5% in boys *vs* 6.9% in girls) and girls at home (55.2% in girls *vs* 33.8% in boys). Boys and girls also showed significant differences (p = 0.001) in the meters of precipitation: girls fell from a median value of 5 m whereas boys fell from a median value of 2 m. Finally, we found noticeable differences in the age of children according to the trauma site: whereas the median age of falls occurring at home was 3 years in boys and 3.5 years in girls, the median age of falls on public roads and during leisure activities was 11 years in boys and 10 years in girls. These age differences were statistically significant only in boys (p < 0.001). When we examined the age and sex of the children that fell from more than 2 stories (6 m), the height associated with severe damage, we found 4 boys and 10 girls of a total sample of 59 cases (38 boys and 21 girls) that reported the meters of precipitation in the database.

The proportion of pedestrian traffic collisions was very similar between sexes (14% in boys, 17% in girls) but boys showed a median age (5 years, IQ 3-8) significantly lower (p = 0.012) than girls (10.5 years, IQ 6-13.7). In both sexes, cars were the principal vehicle involved in the pedestrian traffic collision, which happened mostly on urban roads without established pedestrian crossings.

**Table 2.** Comparison of children’s demographic characteristics in falls. P value indicates the results of sex comparisons.

| Sample characteristics in falls | Boys | Girls | p value |
| --- | --- | --- | --- |
| **Number of cases, *n* (%)** | 90 (72) | 35 (28) | 0.038 |
| **Age** |  |  |  |
| Median (IQ25 - IQ75) | 8 (3-13) | 8 (2 – 11) | 0.350 |
| **Age categories *n* (%)** |  |  |  |
| Neonate | 5 (5.6) | 2 (5.7) | 0.099 |
| Infant | 11 (12.2) | 6 (17.1) |  |
| Young child | 24 (26.7) | 7 (20) |  |
| Child | 21 (23.3) | 13 (37.1) |  |
| Adolescence | 29 (32.2) | 7 (20) |  |
| **Accident site *n* (%)** |  |  |  |
| At home (inside - outside) | 27 (33.8) | 16 (55.2) | 0.0002 |
| Public road | 21 (26.3) | 6 (20.7) |  |
| Leisure<sup>a</sup> | 22 (27.5) | 2 (6.9) |  |
| Other<sup>b</sup> | 10 (12.5) | 5 (17.2) |  |
| **Meters of precipitation** | N = 38 | N = 21 |  |
| Median (IQ25-75) | 2 (1-3) | 5 (3-8) | 0.001 |
| **Age according to accident site, Median (IQ25 - IQ75)** |  |  |  |
| At home (inside – outside) | 3 (0-5) | 3.5 (0-11.25) | 0.499 |
| Public road, Leisure and Other | 11 (4.5-13.5) | 10 (4-11.5) | 0.278 |

<sup>a</sup>Leisure includes outdoor activities, recreational areas, sport areas and rural or natural areas;
<sup>b</sup>Other includes commercial and service areas, other places not recorded and unknown information.

Severity scores and affected body regions are recorded in Table 3. When patients were admitted through a PPT code, priority 2 was the most activated code in both sexes (52.5%). Cases of priority 0 were also frequent (31.7%). Girls showed higher frequencies of PPT 2 (61.1% in girls *vs* 47.7% in boys) corresponding to stable patients with no apparent anatomical injuries but that have suffered a high energy trauma. Boys showed a relatively high frequency of PPT 0 activations (36.9% *vs* 22.2%) involving cases with physiological and clinical characteristics associated with a higher severity. The median values of the Glasgow Coma Scale and the Pediatric Trauma Score were the same in both sexes. The most frequently injured body region was the head (62.9%), followed by the abdomen (17.1%) and the extremities and spine (9.1%). No sex differences were found for any variable of Table 3.

**Table 3.** Severity scores and affected body regions. P value indicates the results of sex comparisons.

| Severity scores | Total | Boys | Girls | p value |
| --- | --- | --- | --- | --- |
| **PPT priority, *n* (%)** | N = 100 | N = 65 | N = 35 |  |
| PPT 0 | 32 (32) | 24 (36.9) | 8 (22.8) | 0.299 |
| PPT 1 | 15 (15) | 10 (15.4) | 5 (14.3) |  |
| PPT 2 | 53 (53) | 31 (47.7) | 22 (62.9) |  |
| **Glasgow Coma Scale (GCS)** | N = 192 | N = 126 | N = 66 |  |
| Median (IQ25-IQ75) | 15 (13 - 15) | 15 (12 - 15) | 15 (14 - 15) | 0.197 |
| GCS categories, *n* (%) |  |  |  |  |
| ≤ 8 | 30 (15.6) | 22 (17.5) | 8 (10.5) | 0.109 |
| 9-13 | 19 (9.9) | 15 (11.9) | 4 (6.1) |  |
| 14-15 | 143 (74.5) | 89 (70.6) | 54 (81.8) |  |
| **Pediatric Trauma Score (PTS)** | N = 187 | N = 122 | N = 65 |  |
| Median (IQ25-IQ75) | 10 (8-11) | 10 (8-11) | 10 (8-11) | 0.983 |
| PTS categories, *n* (%) |  |  |  |  |
| 0-8 | 59 (31.5) | 37 (30.3) | 22 (33.9) | 0.596 |
| ≥ 9 | 128 (68.5) | 85 (69.7) | 43 (66.1) |  |
| **Affected body region, *n* (%)** | N = 175 | N = 118 | N = 57 |  |
| Head | 110 (62.9) | 76 (64.4) | 34 (59.6) | 0.548<sup>a</sup> |
| Chest | 8 (4.6) | 5 (4.2) | 3 (5.3) |  |
| Abdomen | 30 (17.1) | 22 (18.6) | 8 (14) |  |
| Extremities and spine | 16 (9.1) | 9 (7.6) | 7 (12.3) |  |
| Drowning | 5 (2.9) | 2 (1.7) | 3 (5.3) |  |
| Other | 6 (3.4) | 4 (3.4) | 2 (3.5) |  |

<sup>a</sup> Drowning and other have been grouped for the Chi-square test.

Only four children died, representing 1.3% (2/151) of the boys and 2.6% (2/76) of the girls admitted to the pediatric intensive care unit. These cases included two boys (1 year and 5 years old) involved respectively in a traffic collision suffering spinal cord injuries and one drowning, and two girls (1 year and 4 years old) with brain injury, one due to crushing and the other due to a fall.

When, through a bivariate logistic regression, children with severe trauma were compared with other non-trauma PICU patients, sex and age were significant risk factors for severe trauma (Table 4). Boys showed higher odds (OR = 1.59, 95% CI 1.19 – 2.13, p = 0. 002). As age increased so did the risk, with adolescents being the group with the highest odds (OR = 3.77, 95% CI 1.69 - 8.40, p = 0.001).

**Table 4.** Binary logistic regression for severe trauma.

| Logistic regression | OR | 95% CI | p value |
| --- | --- | --- | --- |
| Male sex | 1.593 | 1.189 - 2.134 | 0.002 |
| Age<sup>a</sup> |  |  |  |
| Infant | 0.720 | 0.312 - 1.662 | 0.442 |
| Young child | 3.207 | 1.437 - 7.156 | 0.004 |
| Child | 3.332 | 1.484 - 7.483 | 0.004 |
| Adolescence | 3.770 | 1.691 - 8.405 | 0.001 |
| Nagelkerke's R<sup>2</sup> | 0.085 |  |  |
| Hosmer-Lemeshow test probability | 0.695 |  |  |

<sup>a</sup>Neonate was used as a reference category

## DISCUSSION

In this study we have analyzed the demographic characteristics of a sample of children with severe trauma to determine the type of events causing these injuries, and to measure the contribution of sex and age in each type of injury. A previous study published in 2015 in the same pediatric population indicated that boys suffered more severe trauma events than girls, but we were not able to stratify those injuries into specific causes by sex and age [15]. Thanks to the Hospital Sant Joan de Déu’s participation in a trauma registry (TraumCat) that collects hospital based information of severe trauma patients in Catalonia (Spain), we have had access to a detailed database of pediatric patients. Since this hospital is a reference center, the patients recorded, although the number could seem low (227 patients), are a fairly good representation of the severe injuries of the pediatric population of Catalonia.

A limitation of this study is the fact that *in situ* deaths are not recorded in our database. Thus, our results should be interpreted in a context of children admitted to the hospital alive. As a general frame, in 2014, external causes (including severe trauma) were among the main causes of child death in Catalonia [24]. More concretely, death rates due to all kind of traffic fatalities ranged from 2.7 and 2.8 (per 100,000 boys and girls, respectively) in infants of less than 1 year to 0.24 and 0.52 per 100,000 in boys and girls from 5-14 years old. Death rates from traffic fatalities were higher than death rates by falls in all age categories except in girls from 1-4 years old (death rate 0.65 in falls *vs* 0 in traffic collisions), and in boys from 5-14 years old (death rate 0.49 in falls *vs* 0.24 in traffic collisions).

In our study, falls are the most common type of trauma. Pedestrian and other traffic collisions are far less prevalent. The efforts made in improving road safety, coupled with the fact that traffic collisions can be prevented through the use of seat belts, child safety seats, and airbags may explain the low frequency of children affected. In fact, Spain is cited in the Road Safety Annual Report 2015 of the OECD countries as the country showing the greatest reduction (70%) of traffic collisions between 2000 and 2013 [25].

Comparable studies in other developed countries show road traffic collisions as the main cause of severe trauma although protection measures in these countries are well implemented. Countries such as Canada show motor vehicle crashes (39%) as the most common mechanism of injury, followed by being run over (21%) [26]. France also exhibit a high proportion of motor vehicle crashes (85%) followed by falls (13%) [27], a similar pattern has been described in Croatia with 81.7 % of traffic collisions (being run over included) [28]. In developing countries, more traffic collisions and incidents of being run over are observed due to low passenger protection and a general lack of measures to reduce vehicle crashes [29].

The head has been the most affected body region in children, possibly because of the high number of falls. Our data are in line with previous studies showing that falls are highly related to traumatic brain injury [30,31]. Falls activated the most severe priorities (PPT 0 and 1) in 55.6% of cases, followed by traffic collisions (27.8%) and being run over (16.7%). Boys activated more cases of PPT 0 than girls (36.9% *vs* 22.2%) but the Glasgow Coma Scale, the score recording the consciousness state, was the same in both sexes suggesting that although boys have a higher number of severe trauma events, these events do not put them in a worse clinical situation.

The high frequency of falls in our study, 60.7% of the total observed injury events, is a worrying but not surprising result. Although they are less fatal than in the elderly, falls can be an important cause of severe injury in children, especially when they are from heights greater than 2 stories (6-7 m). Sex, age, and socio-economic status have been reported as independent risk factors. In different studies, the number of boys exceeds those of girls in proportions that ranged from 1.5:1 to 2.1:1. When race or ethnicity is reported, USA Latino and African-American children are overrepresented as a result of living in multiple-story housings, very often in low-income urban neighborhoods [32–34].

A large proportion of falls in our study is observed in boys (72% *vs* 28% in girls) and occurs predominantly on public roads and during leisure activities whereas in girls these are mainly observed at home. Interestingly, the median age of falls is the same in both sexes but shows noticeable differences between falls occurring at home and those observed on public roads and during leisure activities. Falls in this latter category show a median age of 11 years in boys and 10 years in girls, whereas the median age of falls at home is observed at much younger ages (3 years in boys and 3.5 years in girls). Trauma events in public spaces are mainly concentrated in sport and playground areas whereas at home, furniture-related falls and falling down stairs are the most common injuries [34]. The fact that, in our study, half of the children who fell at home were younger than 3 years can be considered the natural result of both their curiosity to explore their close environment and their immaturity in motor and cognitive skills [35]. However, we should keep in mind that these children suffered a fall of enough severity to activate a PPT code of the emergency medical service or to be admitted to a pediatric intensive care unit. Single parenthood, large family size, low parental education and other socio-economic variables related with poverty have been associated with this kind of child injuries [1]. Unfortunately, our database does not allow us to explore any of these variables.

We have found a relatively high representation of children younger than 5 years old, particularly girls, falling from extremely dangerous heights, but this result should be interpreted with caution because it represents a small part of the total falls. In any case, our findings can serve as a warning to parents and authorities to improve safety measures at home, mainly in windows and balconies. This is of particular interest in Barcelona and its metropolitan area, with a large representation of multiple-story housing.

The large proportion of boys observed in falls has not been detected in any of the other trauma events recorded in our database except in pedestrian traffic collisions, where the percentage of boys (61% *vs* 39 % in girls) also points out that differences in risky behavior between sexes may explain the disproportion observed. This tendency could also explain the remarkable sex differences in the age at which pedestrian traffic collisions occurred in our sample (median of 5 years in boys *vs* 10.5 in girls). In fact, we have found that sex and age are statistically significant predictors of severe trauma. Our results can be explained by cultural factors causing boys to be more likely to engage in dangerous activities, social factors where parents encourage boys to take greater risks while treating girls more cautiously, and finally, cognitive factors. Whereas boys have higher physical activity levels and behave more impulsively, girls tend to learn from experience and approach risky situations more cautiously [35, 36]. On public road, girls’ behavior is more thoughtful and cautious [37] whereas boys are more involved in trauma events that include risky street crossing, unsupervised care, use of electronic devices while walking, and poor usage of safety gear [38].

The relatively low in-hospital mortality observed in our study is a good reflection of the coordination between the emergency medical services activating the PPT code and the hospital staff. However, since every child injury is a tragedy, a deep analysis of the determinants of severe trauma is necessary to reduce child injuries and death. Our results confirm the gender-biased pattern of trauma events: falls are the leading cause of injuries, with 2.5 boys for every girl. Falls mostly happened during outdoor leisure activities in boys and at home in girls. The relatively high proportion of young children, particularly girls, involved in falls from great heights needs to be contrasted with data from comparable studies, and if this tendency is confirmed, preventive measures must be launched. A more in-depth analysis trying to determine if socio-economic status is a risk factor in our sample is encouraged.

## ACKNOWLEDGMENTS

Authors thank the PICU medical and nursing staff of the Hospital Sant Joan de Déu, and the working group on TraumCat register. Part of this work has been the final dissertation thesis of RLP. in the Master in Biological Anthropology from the Universities of Barcelona and Autònoma de Barcelona. The authors declare no conflict of interest.

## CONTRIBUTORS

All authors contributed significantly to the paper: SB and RLP recorded data. RLP designed a common database for future exploitations and performed the main analyses. SB, RLP, EE and MEE contributed to the study design, drafted the article and draw the first version of the discussion. LLHP, ME, SP and FJC substantially contributed to the revision of the article. The final version of this work has been seen and approved by all authors. Neither the article nor any part of it have been published or have been submitted for publication elsewhere.

